# Loss of calpain 3 dysregulates store-operated calcium entry and its exercise response in mice

**DOI:** 10.1101/2024.01.12.575391

**Authors:** Katelyn R. Villani, Renjia Zhong, C. Spencer Henley-Beasley, Giorgia Rastelli, Simona Boncompagni, Elisabeth R. Barton, Lan Wei-LaPierre

**Author notes:** Please address correspondence to: Lan Wei-LaPierre, Ph.D. Assistant Professor Applied Physiology & Kinesiology University of Florida 1864 Stadium Rd. Gainesville, FL 32611 Elisabeth R. Barton, Ph.D. Professor Applied Physiology & Kinesiology Associate Dean for Faculty & Staff Affairs College of Health and Human Performance University of Florida 1864 Stadium Rd. Gainesville, FL 32611. These authors contributed equally to this work (Katelyn R. Villani, Renjia Zhong). These authors contributed equal senior author role to this work (Elisabeth R. Barton, Lan Wei-LaPierre).

## Abstract

Limb-Girdle Muscular Dystrophy 2A (LGMD2A) is caused by mutations in the *CAPN3* gene encoding Calpain 3, a skeletal-muscle specific, Ca^2+^-dependent protease. Localization of Calpain 3 within the triad suggests it contributes to Ca^2+^ homeostasis. Through live-cell Ca^2+^ measurements, muscle mechanics, immunofluorescence, and electron microscopy (EM) in *Capn3* deficient (C3KO) and wildtype (WT) mice, we determined if loss of Calpain 3 altered Store-Operated Calcium Entry (SOCE) activity. Direct Ca^2+^ influx measurements revealed loss of *Capn3* elicits elevated resting SOCE and increased resting cytosolic Ca^2+^, supported by high incidence of calcium entry units (CEUs) observed by EM. C3KO and WT mice were subjected to a single bout of treadmill running to elicit SOCE. Within 1HR post-treadmill running, C3KO mice exhibited diminished force production in *extensor digitorum longus* muscles and a greater decay of Ca^2+^ transients in *flexor digitorum brevis* muscle fibers during repetitive stimulation. Striking evidence for impaired exercise-induced SOCE activation in C3KO mice included poor colocalization of key SOCE proteins, stromal-interacting molecule 1 (STIM1) and ORAI1, combined with disappearance of CEUs in C3KO muscles. These results demonstrate that Calpain 3 is a key regulator of SOCE in skeletal muscle and identify SOCE dysregulation as a contributing factor to LGMD2A pathology.

## INTRODUCTION

Limb-Girdle Muscular Dystrophy 2A (LGMD2A) is a rare neuromuscular disease characterized by progressive and symmetrical weakness of proximal limb muscles, ultimately leading to loss of ambulation in ∼1 in 40,000 individuals (1). LGMD2A is caused by mutations in Calpain 3 (*CAPN3*), a skeletal muscle (SKM) specific, calcium (Ca^2+^)-dependent, cysteine protease (2, 3). Symptom onset typically occurs in the second decade of life and includes presentation of waddling gait, tiptoe walking due to Achilles contractures, difficulty with stair climbing, and rising from a seated position (4–7). Males exhibit more severe disease pathology than females, losing ambulation more rapidly (8). However, disease presentation and progression is highly variable, with diagnoses occurring as early as 2 years of age up to mid 60s (2, 4, 9). No genotype-phenotype correlation exists for disease presentation, progression, or severity, and the mechanisms leading to LGMD2A remain poorly understood (10), in part, due to the wide spectrum of mutations throughout the *CAPN3* sequence that alter different properties of the protein (8).

It is well-established that Calpain 3 localizes with and is stabilized by Titin in the contractile apparatus, and it participates in sarcomere remodeling essential to skeletal muscle adaptation through protease cleavage of Titin, among other substrates (6, 11). Disruption of Calpain 3 protease activity through protein instability or mutation with the protease domain was long held to be a primary cause of LGMD2A (12–14) . However, in addition to its Ca^2+^ dependent protease activity, Calpain 3 has also been implicated as a scaffolding protein, where it may help to stabilize complexes in skeletal muscle. Of note is localization of Calpain 3 at the triad (15), a membrane system consisting of one transverse tubule (T-tubule) flanked by two terminal cisternae, the enlarged terminals of sarcoplasmic reticulum (SR). The triad is the primary site for maintenance of Ca^2+^ homeostasis during excitation-contraction (EC) coupling, supported by two key proteins, the voltage sensor Dihydropyridine receptor (DHPR) in the T-tubule membrane, and the Ryanodine receptor (RyR), Ca^2+^ release channels in the SR membrane (16). Previous studies have shown that Calpain 3 co-localizes with the RyR Ca^2+^ release channels as well as the sarcoplasmic/endoplasmic reticulum (S/ER) Ca^2+^ ATPase (SERCA) where it could play a non-proteolytic, structural role (15, 17). However, evidence for how Calpain 3 regulates Ca^2+^ handling is mixed. In *Capn3^-/-^*mice, reduced RyR protein levels were associated with less Ca^2+^ release from the SR upon stimulation, even though Ca^2+^ reuptake via SERCA was not impaired (15, 18). In contrast, myotubes derived from LGMD2A patients displayed downregulation in expression and function of SERCA compared to myotubes from healthy control subjects, suggesting that Ca^2+^ reuptake following muscle activity may be reduced in patients (19). Thus, while there are indications that Calpain 3 contributes to Ca^2+^ homeostasis, its precise role(s) in skeletal muscle Ca^2+^ handling has not been fully defined, nor how the consequences of mutations in *CAPN3* may contribute to LGMD2A pathology.

One consequence of reduced Ca^2+^ reuptake through SERCA is diminished intracellular SR Ca^2+^ stores, which is a trigger for Store-operated Calcium Entry (SOCE). SOCE is a a Ca^2+^ refill mechanism that facilitates the influx of extracellular Ca^2+^ into the cytosol and, particularly in skeletal muscle, aids in sustained force generation during prolonged muscle activity and prevention of muscle fatigue (20, 21). In healthy muscle, SOCE is inactive at rest due to repleted SR Ca^2+^ stores. SOCE is activated by depletion of SR Ca^2+^, and it is mediated by stromal-interacting molecule 1 (STIM1) protein located in the SR membrane and ORAI1 channels located in the T-tubule. STIM1 harbors multiple Ca^2+^ binding sites that are sensitive to SR Ca^2+^ levels. Upon decreased SR Ca^2+^, STIM1 loses its bound Ca^2+^ and undergoes conformational change to form oligomers in the SR membrane. The STIM1 complex interacts with ORAI1 proteins in the T-tubule resulting in the formation of ORAI1 channels that facilitate extracellular Ca^2+^ influx. The STIM1/ORAI1 structures have been termed Ca^2+^ Entry Units (CEU). STIM1 and ORAI1 interact either at the triad junction close to the RyR to facilitate ultrafast Ca^2+^ entry (22) or through the CEUs in the I-band of the muscle contractile apparatus to support sustained Ca^2+^ influx and enable more efficient cross-bridge activation during exercise (20). In response to prolonged exercise, the T-tubule extends into the I-band and forms junctions with remodeled SR stacks of flat cisternae to increase CEU incidence and further enhance Ca^2+^ entry (23). CEU formation and enhanced SOCE are observed in mouse skeletal muscles following 1 hour treadmill running, with SOCE activity returned to baseline within 6 hours after exercise (23). Genetic manipulation leading to massive depletion of SR Ca^2+^ stores can also trigger CEU formation, as observed in calsequestrin1 knockout mice (24). Chronic SOCE activity is detrimental to cell and mitochondrial health and impairs Ca^2+^-dependent signaling pathways due to cytosolic Ca^2+^ overload. Aberrant SOCE activity has been shown to contribute to the dystrophic phenotype in murine Duchenne Muscular Dystrophy (*mdx)* models and delta-sarcoglycan deficient (*Sgcd^-/-^*) mice; genetic manipulation to reduce STIM and ORAI1 activity ameliorates disease phenotype (25–27).

To date, the role of SOCE as a contributing mechanism to LGMD2A pathology has not been investigated. Given the localization of Calpain 3 at the triad and co-localization with RyR1 and SERCA, we hypothesize that loss of Calpain 3 disrupts Ca^2+^ homeostasis and aberrantly activates SOCE, which could be consequential to the muscle pathology and impaired exercise response in LGMD2A. In this study, we investigated the contribution of SOCE to the LGMD2A pathology in Calpain 3 knockout (C3KO) mice. We revealed that loss of *Capn3* results in elevated SOCE at rest, while SOCE activation is reduced in response to treadmill running in C3KO mice, consistent with exercise intolerance in patients with LGMD2A. To our knowledge, this is the first time SOCE dysregulation is documented as a potential mechanism contributing to LGMD2A pathology.

## RESULTS

### Loss of Calpain 3 perturbs calcium handling through chronic SOCE activation

We conducted Ca^2+^ measurements in freshly isolated *flexor digitorum brevis* (FDB) muscle fibers using the ratiometric Ca^2^ indicator Indo-1 to characterize the Ca^2+^ disturbance in C3KO mice. Resting cytosolic Ca^2+^ levels were significantly higher in FDB fibers isolated from 6 month old C3KO mice (Fig. 1A), suggesting intracellular Ca^2+^ overload as a mechanism contributing to the dystrophic pathology in LGMD2A, as it was shown with other types of muscular dystrophies (25–27). This difference was age-dependent, as there were no differences in resting cytosolic Ca^2+^ levels at 8 weeks of age (Indo-1 Ratio_405/485_: C3KO 0.4±0.01; WT 0.4±0.01; mean ± SEM for N=4-6 mice; P=0.94, unpaired t test), paralleling the late-onset and progressive nature of LGMD2A(1). Thus, we extended analysis of Ca^2+^ handling in mice at 6 months of age. Basal Ca^2+^ influx in the absence of pharmacological SR store depletion was significantly increased in C3KO mice compared with WT controls (Fig. 1B), indicating that ablation of *Capn3* elicited constitutively active SOCE. In addition, maximal SOCE measurements following SR store depletion using a depletion cocktail (30 μM Cyclopiazonic acid, 2 μM Thapsigargin and 200 μM EGTA) revealed greater Ca^2+^ influx in C3KO FDB fibers (Fig. 1C & D), confirming enhanced SOCE activity in C3KO mice at rest. This aberrant SOCE in basal conditions may serve as a contributor to increased resting cytosolic Ca^2+^ and LGMD pathology. Calpain 3 was previously identified to colocalize with Ca^2+^-handling proteins including RyR1 and SERCA and loss of *Capn3* reduced levels of these proteins (15, 19). Therefore, we performed western blot analysis for Ca^2+^-handling proteins involved in EC coupling and SOCE to determine if loss of *Capn3* altered their levels. In contrast to previous studies, we observed a significant increase in SERCA1A protein, the fast isoform of SERCA, in quadriceps muscles from C3KO mice (Fig. 1E). STIM1L and STIM1S (two splicing variants of STIM1 involved in SOCE), DHPR, and RyR levels did not differ between genotypes. Gene expression was also examined, including *Orai1,* which was significantly increased with loss of *Capn3*, but no alteration in *Stim1* or *Atp2a1* expression was observed (Fig. 1F). Taken together, the heightened resting SOCE activity and perturbation of Ca^2+^ homeostasis associated with loss of *Capn3* is not due to overt changes in the levels of primary proteins of EC coupling or SOCE.

**Figure 1:**
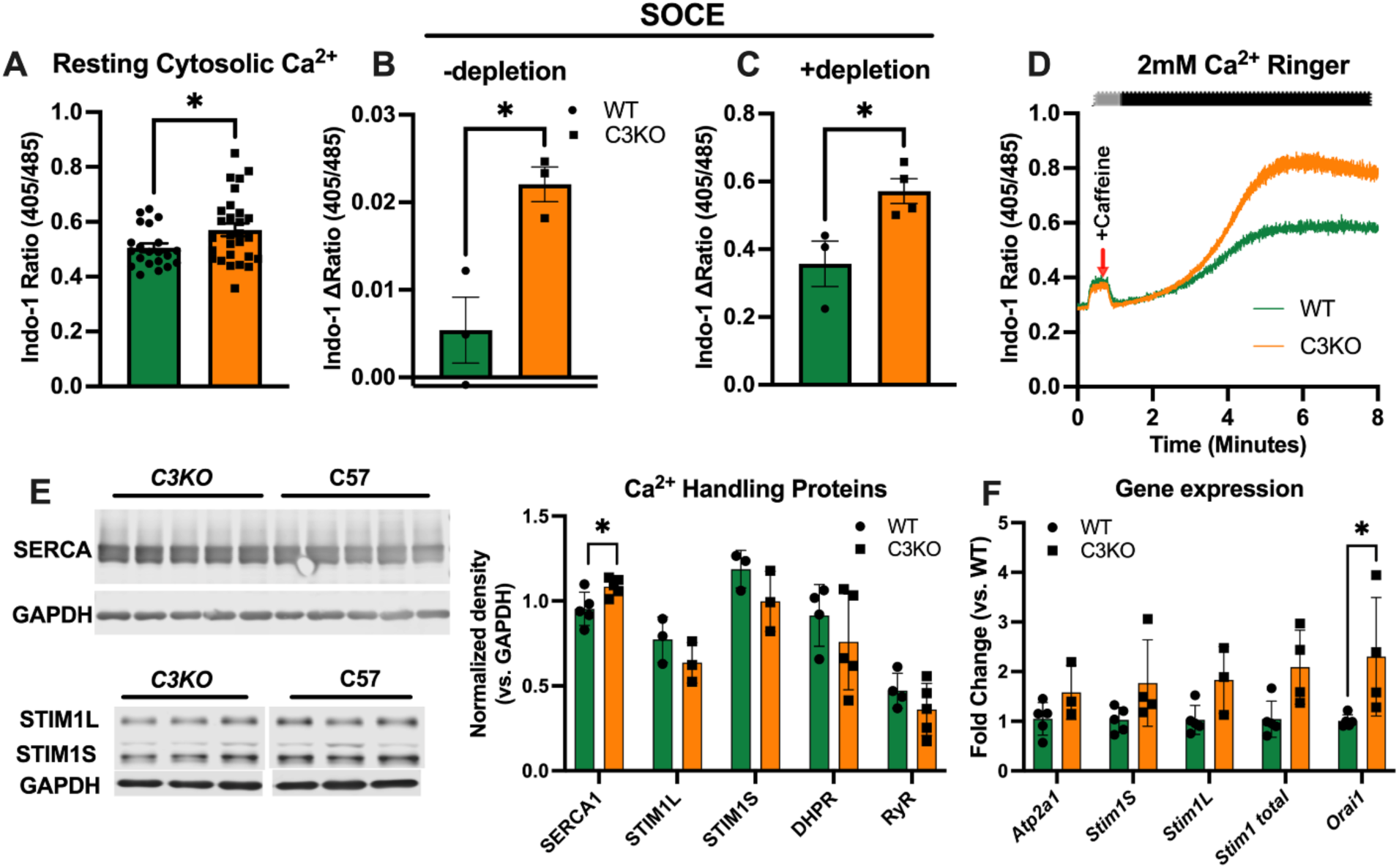
Evaluation of contributors to SOCE and EC coupling in muscles from WT and C3KO mice. (A) Average (±SEM) Indo-1 Ratio_405/485_, indicating resting Calcium levels, in acutely isolated Flexor Digitorum Brevis (FDB) muscle fibers revealed elevated resting cytosolic Ca^2+^. (N=21-28 fibers from 2-3 mice) (B) Average (±SEM) Indo-1 ΔRatio_405/485_ indicating basal SOCE activity without SR store depletion showed aberrantly activated SOCE in C3KO FDB fibers at rest. (N=3-4 mice, 4-6 fibers from each mouse). (C) Average (±SEM) Indo-1 ΔRatio_405/485_ indicating maximal SOCE activity measured in FDB fibers following SR store depletion exhibited greater Ca^2+^ influx with maximal SOCE activation in C3KO muscle cells. (D) Representative traces for maximal SOCE measurements in FDB fibers from WT and C3KO mice in (C). (E) Western blot images and average (±SEM) densitometry quantification of SOCE and EC coupling related proteins. SERCA1A levels were significantly elevated in C3KO quadricep muscle lysates. No differences were observed in STIM1L, STIM1S, DHPR, and RyR levels. (N=3-5 per genotype). (F) qPCR gene expression of SOCE-related genes revealed increased *ORAI1* expression in C3KO quadricep muscles. *Atp2a1* (encoding SERCA1), Stim1S, Stim1L did not differ between genotypes. * P < 0.05 unpaired *Student* t-test, compared with corresponding WT.

### Muscles from C3KO mice are more susceptible to fatigue post-exercise

Previous studies demonstrated that SOCE is activated in mouse skeletal muscles following acute treadmill running, which may help to counteract exercise-induced fatigue (23). Given that many LGMD2A patients suffer from intolerance to exercise (28), disruption of SOCE activation may be an underlying cause. We used a treadmill running protocol as described previously(20) to activate SOCE and assessed force production as an index of SOCE activation following acute exercise in C3KO and WT mice. There were no differences in running duration between strains (WT: 58.6±6.5 min; C3KO: 57.8±5.7 min. mean ± SD for N=12 mice per genotype, P=0.79 by unpaired t-test). *Extensor digitorum longus* (EDL) muscles from non-exercised mice displayed no differences in the pattern of force generation during an *ex vivo* fatigue contraction protocol (Fig. 2A & E). However, C3KO mice showed greater force decay during the fatigue protocol compared to WT EDL muscles within 1-hour post-exercise (Fig. 2B & F), which was evident from the 17th to 38th stimulation. As shown in the forces for the 21^st^ stimulation (Fig. 2D-G), there was a significant decrease in specific force in C3KO EDL muscles 1 hour after treadmill running. This may be attributed to diminished exercise-induced activation of SOCE in C3KO mice. C3KO and WT EDL muscles showed similar fatigability 6 hours after exercise (Fig. 2C & G), when SOCE activity presumably returned to baseline (23). This result indicates that the transient SOCE activation in WT mice that occurs to fight fatigue in response to exercise is lacking in C3KO mice.

**Figure 2:**
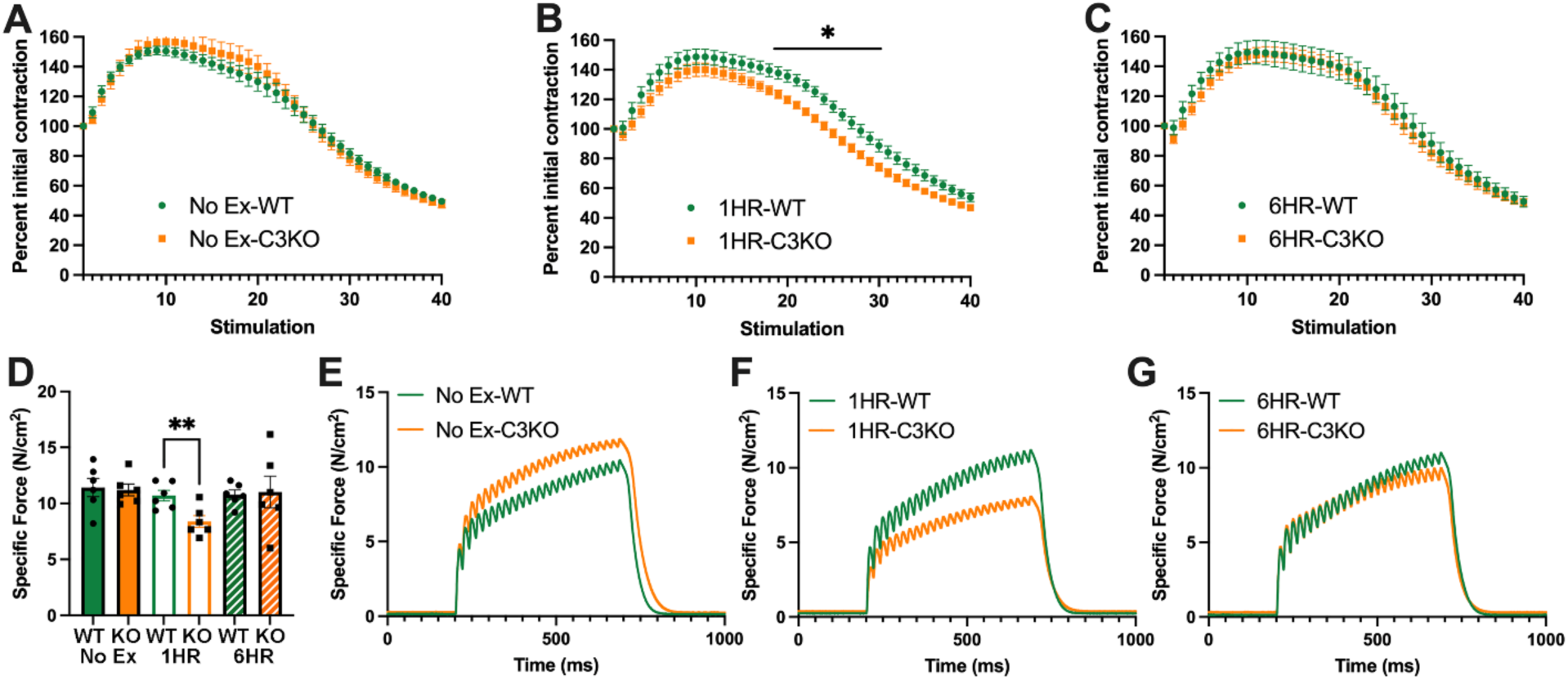
Contractile force measurements during fatiguing stimulation in EDL muscles following treadmill exercise. EDL muscles from WT and C3KO mice were subjected to a fatiguing stimulation protocol (40 consecutive, 500 ms duration, 50 Hz stimulus trains delivered every 2.5 s) at (A) no exercise; (B) 1-hour, when C3KO mice were more susceptible to fatigue; and (C) 6-hours post-treadmill exercise, when the difference in fatiguability in WT and C3KO mice resolved. Fatigue mechanics were analyzed with multiple unpaired *Student* t-tests. (D) Average (±SEM) tetanic force at the 21^st^ stimulation 1HR post-exercise, showing C3KO EDL muscles produce lower force. (E-G) Representative trace of specific forces production, at the 21^st^ stimulus train at (E) no exercise, (F) 1-hour and (G) 6-hour post-exercise. * P < 0.05 compared with corresponding WT at the same time point; unpaired *Student* t-test, N=6 mice per strain, per timepoint.

### Disrupted Ca^2+^ cycling mirrors force production post-exercise in C3KO mice

To determine if the reduced force production in EDL muscles from C3KO mice during repetitive stimulation following treadmill exercise could be attributed to a Ca^2+^ disturbance, we monitored the Ca^2+^ transient decay in FDB fibers. Fibers were from the same cohort of WT and C3KO mice subjected to treadmill running as reported in Fig. 2 at the same time points: no exercise, 1 hour after exercise, and 6 hours after exercise, using a repetitive high-frequency stimulation protocol identical to what was used for EDL muscle force measurements (Fig. 2). Peak Ca^2+^ transients during the stimulation train were detected using mag-fluo-4, a moderate-affinity Ca^2+^ indicator. No significant differences were observed in the Ca^2+^ transient decay profile between FDB fibers from WT and C3KO mice without treadmill exercise (Fig. 3A & E). Consistent with the EDL force measurements, FDB fibers isolated from C3KO mice within 1 hour post-exercise exhibited a greater decline in Ca^2+^ transients during the repetitive stimulation train (Fig. 3B & F), with significant reductions in peak Ca^2+^ transient amplitudes in the second half of consecutive stimulation (stimuli 21–40, #40 stimulus ΔF/F_0_: WT: 0.89±0.03 vs. C3KO: 0.78±0.03) (Fig. 3B & D). The Ca^2+^ transient profile in C3KO FDB fibers paralleled the WT profile 6 hours after exercise (Fig. 3C & G). The changes in peak Ca^2+^ transients mirrored the changes in force observed in EDL muscles, exemplified by the significant decrease in the 21^st^ stimulation 1 hour after exercise (Fig. 3D). In addition, peak Ca^2+^ transient amplitude at the beginning of the simulation train was also reduced in C3KO FDB fibers compared with that in WT mice (Fig. 3B, first stimulation, ΔF/F_0_: WT: 0.90±0.03 vs. C3KO: 0.83±0.03), indicating that a single bout of 65 minutes treadmill running is sufficient to induce a reduction in Ca^2+^ release, possibly due to the lack of SOCE activation leading to store depletion in C3KO mice. These results indicate that disturbed intracellular Ca^2+^ cycling may underlie the reduced fatigue resistance observed in EDL muscles in C3KO mice post-exercise.

**Figure 3:**
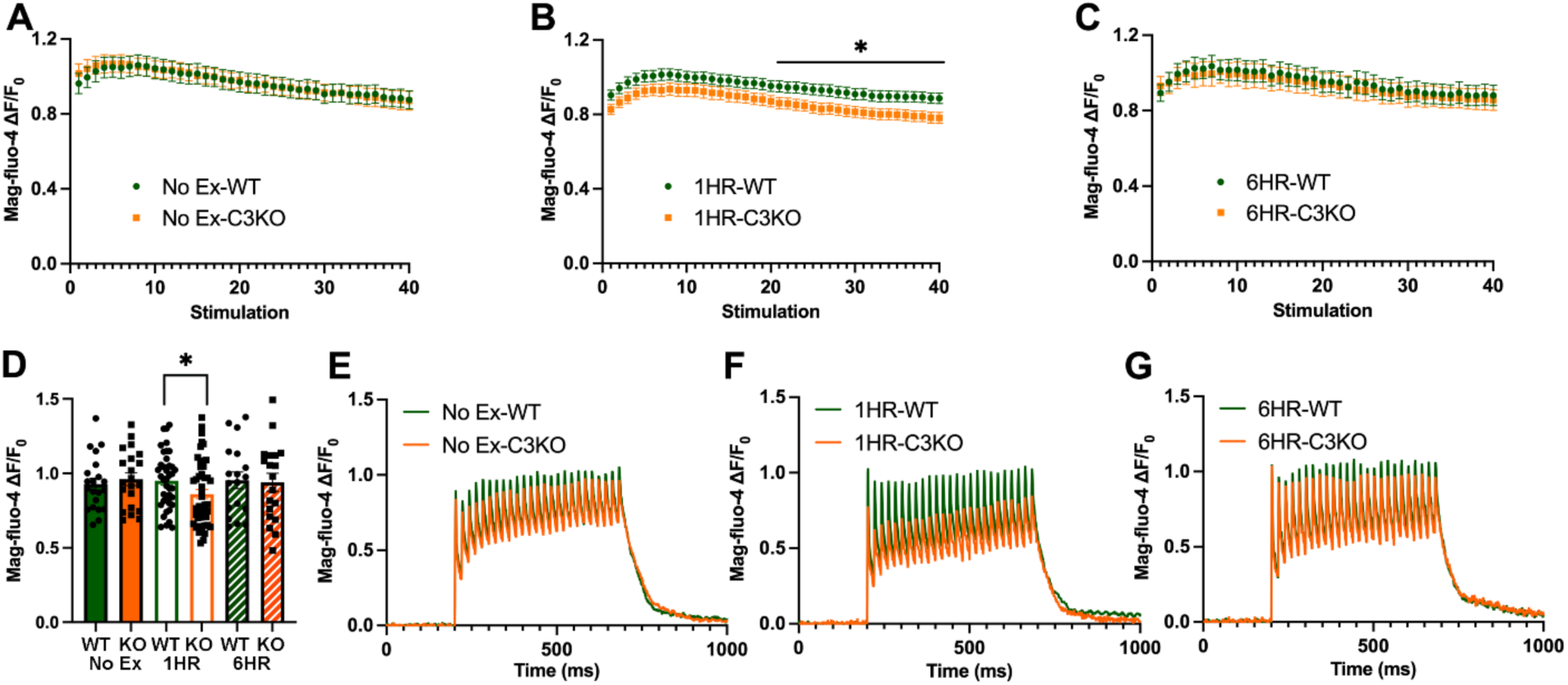
Calcium transient measurements in FDB fibers following treadmill running. Acutely isolated FDB fibers from WT and C3KO mice were subjected to a repetitive tetanic stimulation protocol (40 consecutive, 500 ms duration, 50 Hz stimulus trains delivered every 2.5 s) at (A) no exercise, (B) 1-hour, and (C) 6-hours post-treadmill exercise. Peak Ca^2+^ transient amplitudes during the stimulation train were monitored using Ca^2+^ indicator mag-fluo-4. (A-C) Average (±SEM) ΔF/F_0_ mag-fluo-4 fluorescence in isolated FBD fibers in WT and C3KO revealed less sustained Ca^2+^ transient amplitude in C3KO mice during the repetitive stimulation train at 1-hour (1HR) post-exercise, with comparable decay in transient peak between WT and C3KO at no exercise (A) and 6-hours post-exercise (6HR, C). (D) Average (±SEM) ΔF/F_0_ mag-fluo-4 fluorescence at the 21^st^ stimulation train. (E-F) Representative superimposed mag-fluo-4 (ΔF/F_0_) traces during the 21^st^ stimulation train at (E) no exercise, (F) 1-hour, and (G) 6-hours post-treadmill exercise. * P < 0.05 compared with WT at corresponding time point, unpaired *Student*-t test; n=18-49 fibers from 5-8 mice.

### STIM1 and ORAI1 co-localization increases 1HR post-exercise in WT, but not in C3KO mice

Previous studies showed that co-localization of STIM1 and ORAI1 and the formation of CEUs that support SOCE activation can be visualized by immunofluorescence of muscle fibers (20). To determine if the localization of SOCE proteins differed between WT and C3KO mice in response to exercise, EDL muscle bundles were stained with different combinations of antibodies recognizing STIM1, ORAI1 and RyRs. In WT and C3KO mice at non-exercised conditions, ORAI1 was observed as two peaks in the fluorescence profiles that overlapped with double peaks of RyR (Fig. 4A & B), indicative of its localization in the triad at rest. At 1-hour post-exercise, ORAI1 signals shifted from the triad and co-localized with STIM1 in the I-band in EDL bundles from WT mice, visualized as single, overlapping peaks of ORAI1 and STIM1 in the fluorescence plot (Fig. 4C). The increased ORAI1/STIM1 co-localization was the result of CEU formation and likely was at the basis of increased SOCE. However, this ORAI1 I-band shift was markedly limited in the C3KO mice, where ORAI1 largely remained at the triad with RyR (Fig. 4D). The lack of increased co-localization between ORAI1 and STIM1 upon exercise indicates that CEU formation was hampered in the absence of *Capn3*. At 6-hours post-exercise, STIM1 and ORAI1 no longer co-localized in WT mice (Fig. 4E), indicating exercise-induced SOCE activation resolved, consistent with previous findings (20). In samples from C3KO mice, there was no apparent difference in the shift of ORAI1 (Fig. 4F). More extensive quantification of at least 50 sarcomeres per sample substantiated the significant increase in ORAI1 single peaks in the I band following exercise in WT muscles and the stark absence of this response in C3KO muscles (Fig. 4G). Taken together, the lack of ORAI1/STIM1 co-localization in response to treadmill running, combined with diminished protection of fatigue and reduced Ca^2+^ transients provide compelling evidence for the importance of Calpain 3 in SOCE regulation.

**Figure 4:**
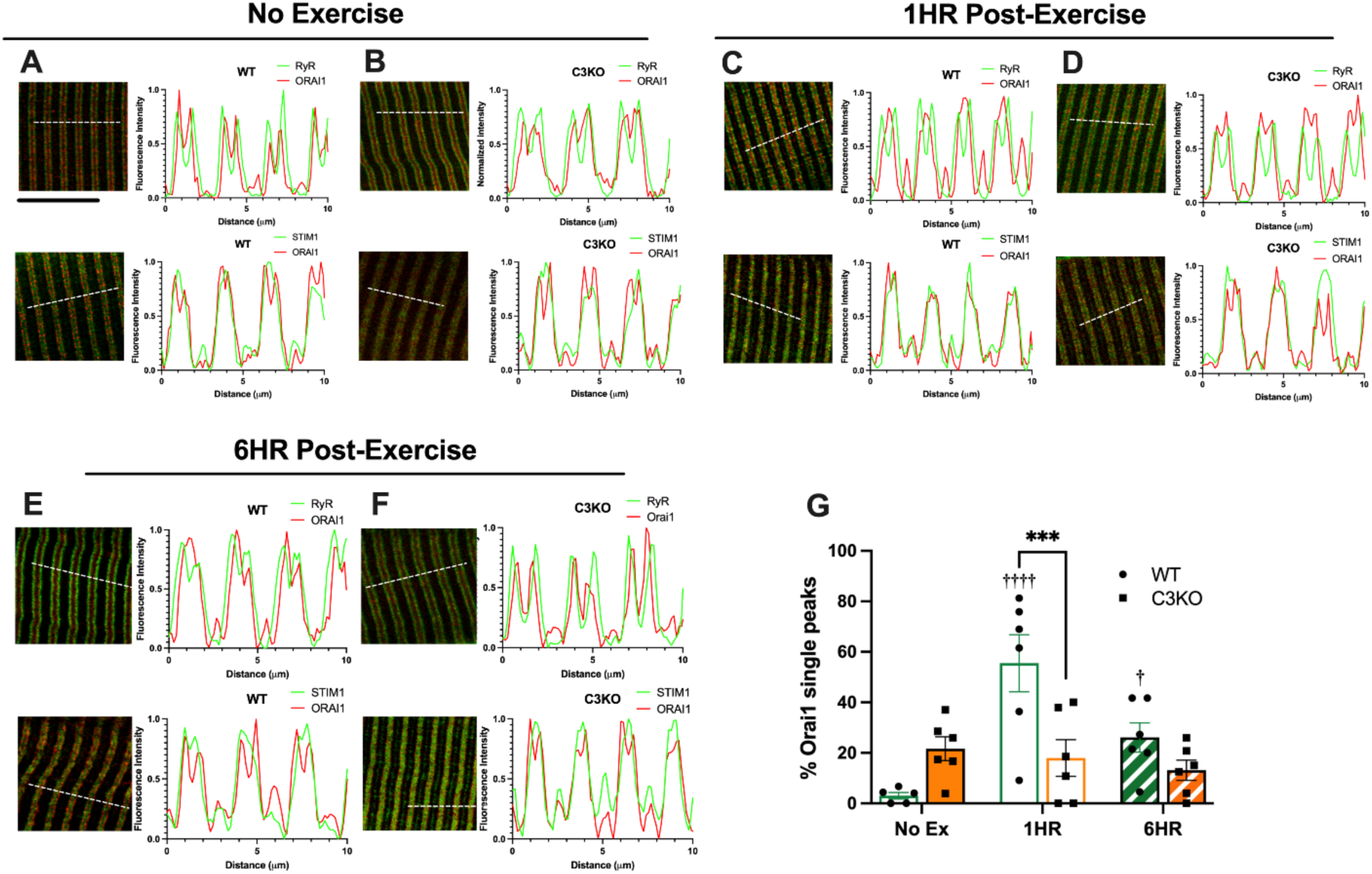
Immunofluorescence analysis on localization of SOCE related proteins. Representative immunostaining images of EDL muscle bundles for SOCE proteins, STIM1, ORAI1, and RyR, with corresponding fluorescence intensity profiles at no exercise (A-B), 1HR (C-D) and 6HRs (E-F) post-exercise. At no exercise, ORAI1 was in the triad and co-localized with RyR in both C3KO and WT mice. 1HR post-exercise, ORAI1 co-localized with STIM1 in WT mice, but not in C3KO mice. The co-localization of ORAI1 and STIM1 resolved at 6HR post-exercise. Fluorescence intensity profiles showing co-localization of proteins across five sarcomeres were plotted using FIJI (ImageJ) software; dashed white line indicative of region plotted. Solid black line in panel A indicates 10 μm. (G) Quantification of the proportion of ORAI1 single peaks per fiber. Mean ±SEM of measurements in one fiber shown Points represent the mean of ∼500 sarcomeres in one fiber (two fiber images from each mouse, N=3 per condition). ***, P<0.01 between genotypes; †, P<0.05, ††††, P<0.001 between conditions within genotype; two-way ANOVA followed by Sidak’s *post-hoc* test.

### C3KO muscles have constitutively assembled CEUs at rest that are disrupted 1 hour after exercise

In healthy murine muscle, exercise induces remodeling in which SR membranes form flat, parallel, multi-layer stacks of cisternae, accompanied by T-tubule elongation toward the Z-line (20). This enables contacts between the extended T-tubule and SR stacks to form CEUs and facilitate influx of extracellular Ca^2+^. To verify a possible correlation between the altered SOCE activity and assembly of CEUs at rest and following exercise in C3KO mice, we utilized EM to quantify the presence of two different components of CEUs, i.e. SR stacks (Fig. 5) and elongated T-tubules (Fig. 6). EDL muscles were from the same cohort of WT and C3KO mice subjected to treadmill running as reported in Fig. 2-4.

**Figure 5:**
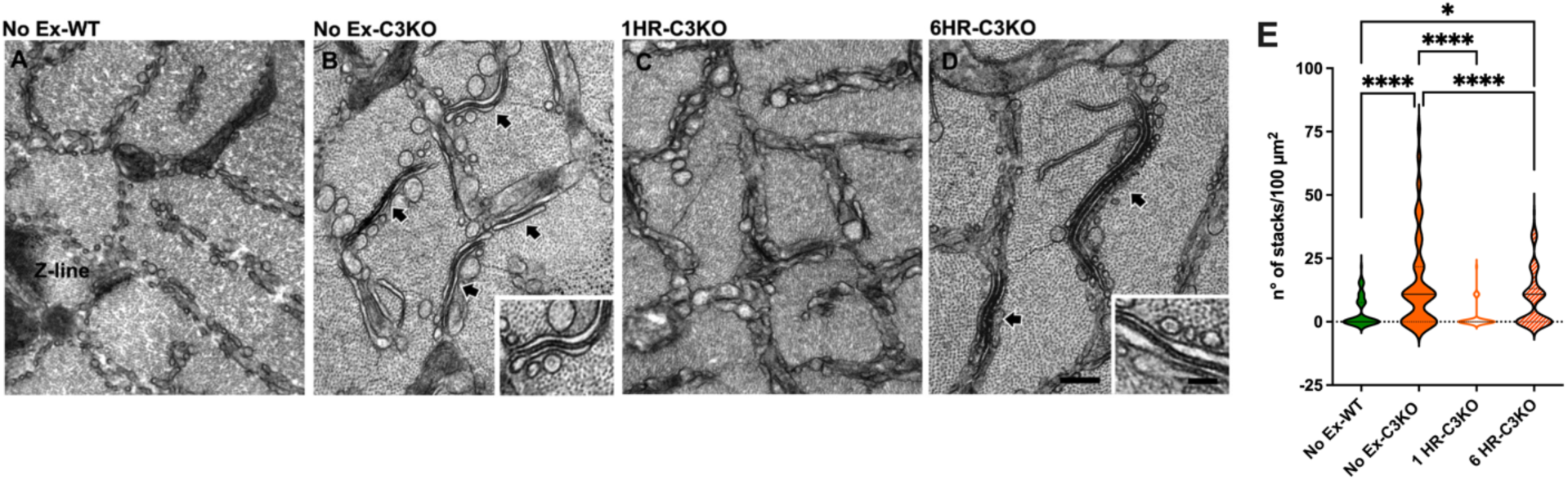
EM analysis of muscles from no exercise, 1 and 6 hours post exercise. Representative EM images of cross sections of EDL muscles from WT no exercise (A), C3KO no exercise (B), C3KO 1HR post exercise (C), and C3KO 6HR post exercise (D). Labeling: large, white-outlined black arrows point to stacks of multiple flat cisternae of membranes (T-tubule and or SR). E. Quantification of SR stacks showed a 3-fold higher number of stacks apparent in no exercise C3KO muscles, the loss of stacks 1HR after exercise, and the re-formation of stacks at 6HR post exercise. Data are shown as violin plots for 75-100 measurements from each muscle sample. Solid black lines indicate medians and dotted lines indicate quartiles. N=3 mice, 1 EDL/mouse for all conditions. Comparisons by 1-way ANOVA and Tukey’s *post-hoc* test; *, P<0.05; ****, P<0.0001.

**Figure 6:**
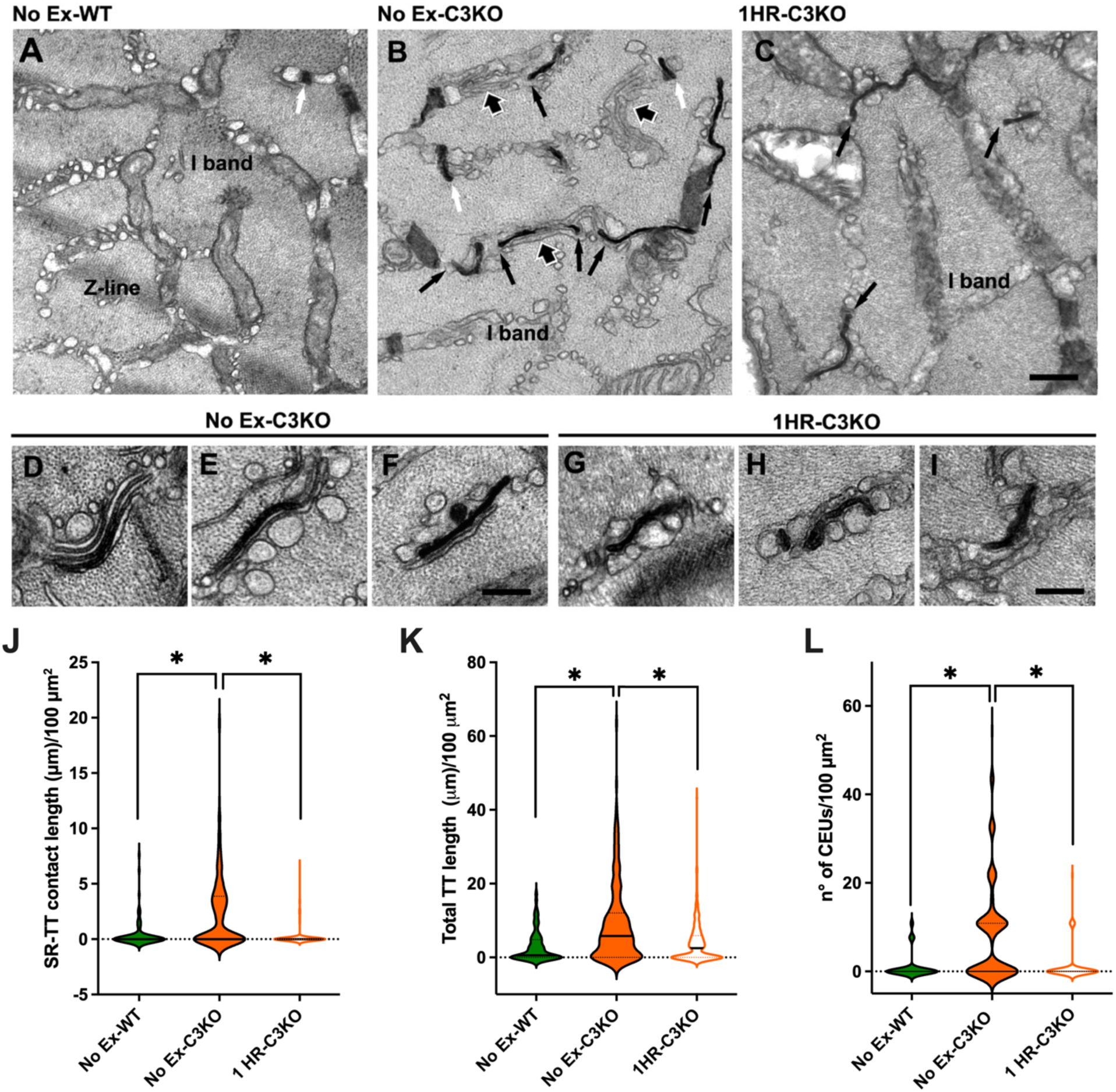
EM analysis of stained T-tubules in muscles from no exercise and 1 hour post exercise. Representative EM images of cross sections of EDL muscles showing T-tubules stained in black with ferrocyanide precipitate from WT no exercise (A), C3KO no exercise (B), and C3KO 1HR post exercise (C). Labeling: white, small arrows in A and B point to T-Tubule of longitudinal triads; black small arrows point to elongated T-tubules; large, white-outlined black arrows point to stacks of multiple flat cisternae of membranes (T-tubule and/or SR). Higher magnification EM images showing SR and T-tubule contacts in no exercise C3KO (D-F) and 1HR post-exercise C3KO (G-I). Scale bars: A-C, 0.5 μm; D-I, 0.2 μm. Quantitative analyses of the T-tubule length at the I band (J) (expressed as μm /100 μm^2^ of cross-sectional area); T-tubule/SR contact length (K) (expressed as μm/100 μm^2^ of cross sectional area); number of CEUs per 100 μm^2^ of cross-sectional area (L). Data for K-L are shown as violin plots for 75-100 measurements from each muscle sample. Solid black lines:medians, dotted lines:quartiles. N=3 mice, 1 EDL/mouse for all conditions. Comparisons by 1-way ANOVA and Tukey’s *post-hoc* test; *, P<0.05; ****, P<0.0001.

As shown in Fig. 5, qualitative examination of EDL fiber cross-sections showed that in muscles from non-exercised WT mice, the I-band SR appeared to be composed of small vesicles arranged into two or three layers between the myofibrils (Fig. 5A). In contrast, in C3KO non-exercised mice, stacks of two (occasionally three) elements of flattened and parallel cisternae of SR membranes were often visible (Fig. 5B). At 1HR post exercise, there was a disruption of the stacks, and instead, the I band SR was mostly comprised of multiple layers of vesicles (Fig. 5C). Vesicles in the 1HR post exercise C3KO muscles were more dilated than those observed in WT non-exercised muscles (Fig. 5A and C, insets). Surprisingly SR stacks were present in C3KO mice 6HR post exercise (Fig. 5D). Quantification of the incidence of SR stacks (defined by at least two SR elements) demonstrated a 4-fold increase in the number of SR-stacks/100 µm^2^ at the I band in muscles from resting C3KO mice compared to that of non-exercised WT mice (Fig. 5E). The number of SR-stacks was significantly lower at 1HR post exercise, but more abundant at 6HR post exercise compared to resting WT muscles (Fig. 5E). Note that at higher magnification small electron-dense strands likely reflecting STIM1 aggregates were visible within the junctional gap between SR/SR membranes (Fig. 5B and D, insets).

In order to enable direct visualization of T-tubule membranes in fibers from non-exercised and 1HR post exercise mice, muscles were stained with Ferrocyanide which forms a dark and electron dense precipitate inside the T-tubule’s lumen (Fig. 6). At rest, muscles from C3KO mice displayed abundant T-tubule extensions in the I band, in contrast to those from WT mice (Fig. 6A, B, J), which afforded greater SR-TT contact (Fig. 6K) and higher incidence of CEUs (Fig. 6L). This observation was consistent with the elevated resting SOCE activity measured in fibers from C3KO mice (Fig.1). The enhanced SOCE at rest is also supported by increased incidence of SR stacks and rounded, vesiculated SR membranes in C3KO muscles (Fig. 6D-F), similar to that reported in WT EDL muscles after treadmill exercise (20). At 1HR post-exercise, the number of CEUs in C3KO mice significantly decreased (Fig. 6C, L), predominantly due to disruption of the assembly of flat stacks of cisternae and the extensive vesiculation of SR membranes (Fig. 6G-I). The predominance of rounded SR morphology not only diminished number of CEUs, and also the SR-T-tubule contact length (Fig. 6K). T-tubule extension also retracted in C3KO EDL muscles following treadmill exercise (Fig. 6J, C3KO 1HR vs. C3KO No execise). Although elongated T-tubules were still observed in the I band in C3KO 1HR post-exercise (Fig. 6C black arrows, G-I), the average length of T-tubule did not reach statistical significance compared with the basal condition (WT no exercise) (Fig. 6J).The lack of CEU formation corresponds to the increase in fatiguability and failure of ORAI1 and STIM1 co-localization post-exercise (Fig. 2-4). Thus, exercise caused a transient loss of CEUs in C3KO muscles, a response that was completely opposite to that found in healthy muscles (20, 23). Taken together, this places Calpain 3 as a central regulator of SOCE through SR stack and CEU formation and stabilization.

## DISCUSSION

Calpain 3 is an atypical member of the calpain family, harboring both protease and scaffolding properties (17, 29). Thus, consequences of *CAPN3* mutations on LGMD2A pathology has been challenging to define. There is general consensus that altered Calpain 3 protease activity and its instability are significant contributors to the disease (13, 30), but whether additional properties of Calpain 3 are critical for skeletal muscle health have not been substantiated. Previous studies have linked Calpain 3 to Ca^2+^ regulation through its interactions with Ca^2+^ handling proteins as well as downstream Ca^2+^ dependent mediators for muscle adaptation (15, 19, 31), but no studies, to date, have examined SOCE in models of LGMD2A. In the current study, we showed loss of Calpain 3: 1) elicited chronic SOCE activation and increased maximum Ca^2+^ influx capacity at resting conditions; 2) induced an increase in cytosolic resting Ca^2+^ levels; 3) dampened SOCE activation in response to exercise; and 4) transiently disrupted SR stack formation following exercise. These findings, while unanticipated, are consistent across multiple complementary measures, and may help to explain key symptoms associated with LGMD2A, including exercise intolerance. We assert that the heightened SOCE activity at rest in muscles of C3KO mice contributes to increased resting cytosolic Ca^2+^, leading to damage of muscle proteins and mitochondria. SOCE activity has been shown to contribute to calcium dysregulation in DMD and sarcoglycanopathies (25, 26), and may be a pathology common to many neuromuscular diseases. However, the reduction of indicators of SOCE coupled with SR vesiculation after exercise in C3KO mice points to additional roles that Calpain 3 plays in Ca^2+^ regulation, and a need to explore these mechanisms to obtain a deeper understanding of LGMD2A pathology.

This is the first time increased resting cytosolic Ca^2+^ has been documented in a LGMD2A mouse model, although intracellular Ca^2+^ overload has been previously shown to promote a dystrophic phenotype in DMD by over activating calpains (25, 26) or triggering mitochondrial membrane permeability transition, leading to mitochondrial damage (32). Indeed, C3KO mice display abundant and disorganized mitochondria, and mitochondrial damage is found in LGMD2A patient biopsies (33–35). As we observed increased cytosolic Ca^2+^ only at 6 months of age, and not at 8 weeks of age, this suggests that there is a slow progression of pathology that leads to this defect, and is consistent with later onset of the disease in humans (4). Examination of C3KO muscles for differences in proteins that directly regulate Ca^2+^ in this and other studies have not reached consensus on one clear culprit for disrupted Ca^2+^ homeostasis (36, 37). In our hands, increased *Orai1* transcript expression supported the possibility of elevated resting SOCE, even though STIM1 levels did not change. Although an increase in *Orai1* transcript was detected, elevation in ORAI1 protein expression could not be confirmed due to the lack of specific ORAI1 antibodies that can be reliably used for immunoblotting. Even so, therapeutic reduction of resting SOCE in LGMD2A muscles, irrespective of the detailed mechanism, may benefit many individuals with this disease and slow progression of pathology.

Dysregulation of SOCE activation post-exercise with ablation of *Capn3* could shed light on the mechanism for the exercise intolerance in LGMD2A patients (38). There are multiple lines of evidence supporting limited exercise-induced SOCE activation in C3KO mice: increased fatigability in EDL muscles, more rapid Ca^2+^ transient decay in FDB fibers during repetitive electrical stimulation, poor localization between STIM1 and ORAI1 in C3KO muscles at 1-hour after treadmill exercise, which is accompanied by disruption of flattened SR stacks. Immunostaining of EDL muscle bundles in WT mice confirmed the findings in previous studies (20, 23) on ORAI1 translocation into the I-band and co-localizing with STIM1 post-exercise, supporting formation of CEU and increased SOCE. Limited co-localization between ORAI1 and STIM1 in C3KO mice along with loss of CEUs apparent by EM suggests that there is an exercise-dependent instability of stack formation. How *Capn3* deficiency impairs muscle remodeling following exercise is unclear. If Calpain 3 protease activity were important for stack and CEU formation, then in its absence there would be limited CEUs, and SOCE would be diminished throughout rest and following exercise. Our observations of heightened resting SOCE argue against this. An alternative possibility is that Calpain 3 serves as a scaffolding protein interacting with titin as well as additional proteins involved in contact between T-tubule and SR stacks. The disruption of SR stacks points to the need for Calpain 3 to serve as part of a scaffold that enables the cisternae to remain flattened. The requirement of Calpain 3 for T-tubule extension and remodeling is uncertain. T-tubule extension occurs in the absence of Calpain 3 at rest. However, this T-tubule remodelling becomes limited in C3KO mice following exercise. Taken together, Calpain 3 influences T-tubule elongation and SR remodeling, and the mechanisms by which this occurs warrants further investigation. Because our findings are the consequence of total ablation of *Capn3*, these may not accurately replicate patient pathology in which mutations are widely dispersed throughout the *CAPN3* gene. Approximately one-third of patients produce a full-length protein that retains proteolytic activity (39) and other documented mutations specifically impair Calpain 3’s ability to bind to titin, yet still cleave titin (40). How or if these specific mutations impact SOCE activity and muscle fatiguability or yield different consequences is unknown. Future examination of different *CAPN3* mutations with its proteolytic function/titin association either abolished or preserved is essential to understand the contribution of Calpain 3 in SOCE activation, Ca^2+^ homeostasis regulation and the mechanism for exercise intolerance in LGMD2A.

In summary, we have demonstrated for the first time that aberrant SOCE activation and intracellular Ca^2+^ overload may contribute to the muscle pathology with loss of *Capn3*. Further, attenuated formation of CEUs in response to exercise potentially serves as a mechanism for exercise intolerance in *Capn3* deficient mice. Identifying SOCE as a novel mechanism for LGMD2A provides new therapeutic strategies for a disease in which no therapies are currently available for clinical use.

## METHODS

### Animal studies

All procedures conducted in the study are in accordance with the guidelines of Institutional Animal Care and Use Committee and approved by the University of Florida. C57BL/6J mice and previously generated C3KO male mice were used for this study (41). Only males were used to reduce sex related variability of disease phenotype, which is most evident in male mice at 6 months of age. All mice were housed in the animal facility with a 12-hour light and 12-hour dark cycle with *ad libitum* access to food and water.

### Treadmill Protocol

A treadmill protocol was performed at room temperature using a running treadmill (Columbus Instruments, Columbus, OH, USA) on a flat surface (0° incline) as previously described (23). The protocol included a warm-up period of 10 min at 5 m/min. The exercise protocol began with an initial 25 min at a speed of 10 m/min, followed by 20 min at 15/m/min, 15 min at 20 m/min and then five final 1 min intervals where the speed was increased an additional 1 m/min for each interval. Motivation to continue running was done with light tapping with forceps to their rumps. The protocol was stopped when mice either reach the end of the protocol or were unable to continue as indicated by the inability of the animal to maintain running, indicated by 50 cumulative taps. Following treadmill exercise, *extensor digitorum longus* (EDL) muscles (for *ex vivo* muscle contractility studies) and *flexor digitorum brevis* (FDB) muscles (for Ca^2+^ measurements) were removed and appropriately prepared for experiments at no treadmill exercise (control), 1-hour post treadmill-exercise, and 6-hour post-treadmill exercise time points.

### *Ex Vivo* EDL muscle mechanics

For muscle fatigue evaluation, mice were anesthetized with a combination of ketamine (10 mg/kg) and xylazine (80 mg/kg) diluted in saline. Upon anesthetization at no exercise, 1-hour and 6-hours post-treadmill exercise, EDL muscles were dissected and placed in a bath of gas-equilibrated (95% O_2_/ 5% CO_2_) Ringers solution (120mM NaCl, 4.7mM KCl, 2.5mM CaCl2, 1.2mM KH2PO4, 1.2mM MgSO4, 25mM HEPES and 5.5mM Glucose) maintained at 37°C. The proximal tendon was sutured to a rigid hook and the distal tendon was tied with 6-0 silk suture to a servomotor arm (Aurora Scientific, Ontario, CAN), which served as the force transducer. Stimulation was delivered via two platinum plate electrodes positioned along the length of the muscles. Stimulation parameters were controlled by Aurora developed software and delivered via a bi-phase current stimulator (Aurora Scientific, Ontario, CAN). Optimum length (L_o_) of the muscle was defined as the length at which maximal twitch force develops from supramaximal stimulation. After establishing L_o_, muscles were equilibrated using three tetani for a train of pulses for 500 msec at 150 Hz, given at 1 min intervals. After a 1 min of rest, muscles were subjected to a repetitive stimulation protocol (40 consecutive, 500 ms duration, 50 Hz stimulus trains delivered every 2.5 s). Muscle specific forces (N/cm^2^) were calculated by normalizing muscle tension to the muscle cross-sectional area (CSA). Physiological CSA was estimated using the following formula: CSA = muscle mass (g)/[L_o_ (cm) x (L/L_o_) x1.06 (g/cm^2^)], where L/L_o_ is the fiber length to muscle length ratio (0.45 for EDL), and 1.06 is the density of muscle.

### Gene expression

Transcripts of genes encoding Ca^2+^ handling proteins were quantified using quantitative PCR (qPCR). Total RNA from skeletal muscle was isolated using TRIzol reagent (Life Technologies) according to the manufacture protocol and treated with recombinant Rnase-free DNAse I (Thermo Scientific). RNA concentration and purity were determined with NanoDrop 2000 Spectrophotometer (ND-2000, Thermo Fisher Scientific). RNA (1 mg) from each sample was subjected to reverse transcription with High capacity cDNA kit (Applied Biosystems). Duplicates of the resultant cDNA were used for real time PCR using SYBR Master Mix (Applied Biosystems) and oligo primers for target genes and carried out on QuantStudio 3 (Applied Biosystems). Transcripts of SOCE and Ca^2+^ signaling related proteins were quantified in quadriceps muscles from 6 months old WT and C3KO mice. All data were normalized to 18S as a housekeeping gene and plotted as Log_2_ fold changes. Primer sequences for each transcript analyzed are presented in Table 1.

**Table 1:**
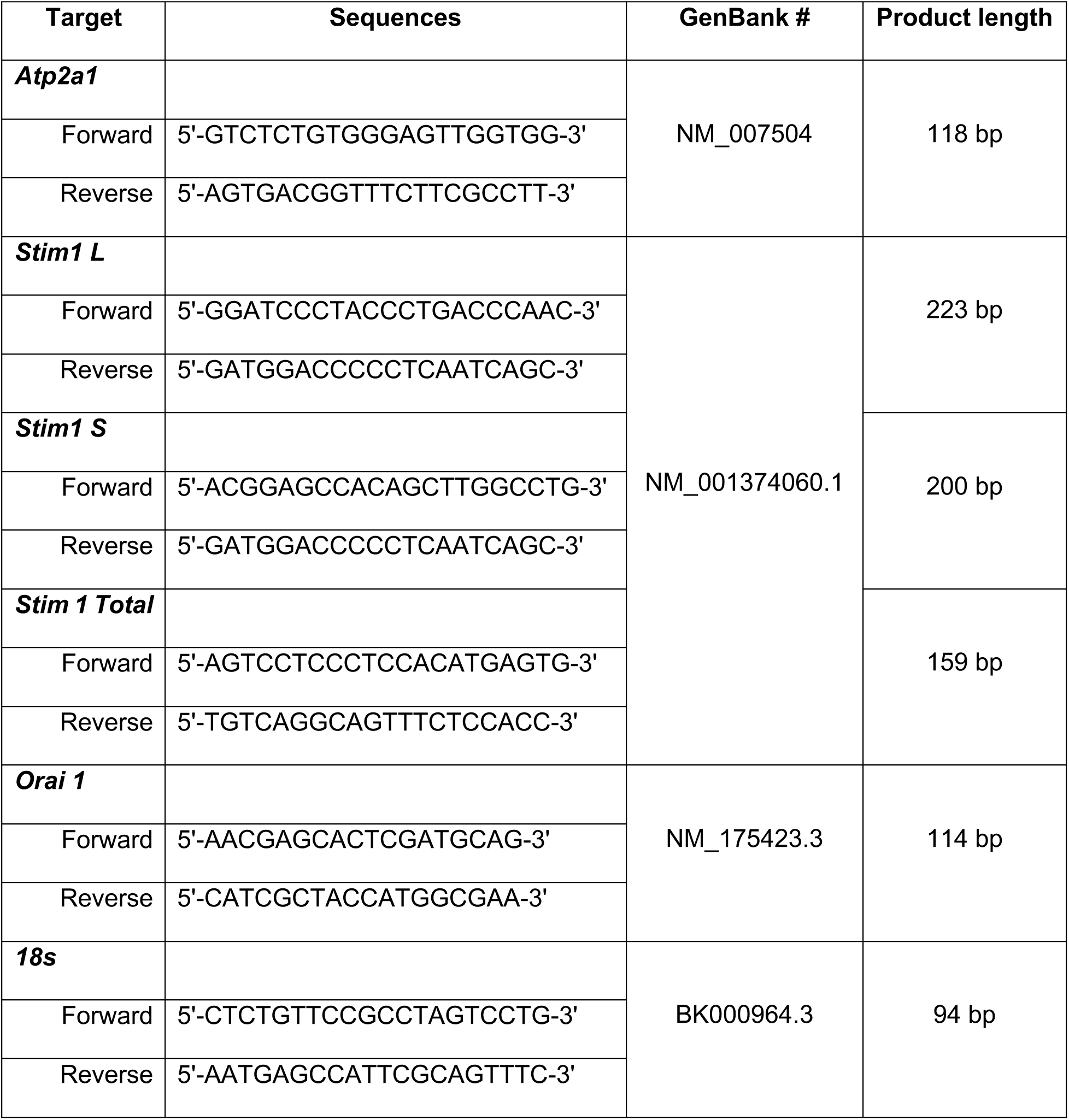
Primer sequences used for quantitative PCR.

### Immunoblotting

The overall levels of Ca^2+^ signaling related proteins (RyR, DHPR, SERCA, STIM1L and STIM1S were quantified using SDS-PAGE followed by immunoblotting in Quadriceps muscles in 6 months old WT and C3KO mice. Tissues extracted for immunoblot analysis were snap-frozen in liquid nitrogen and stored at -80°C until further processing. Quadriceps muscles were mechanically ground by Mortar and Pestle in dry ice and homogenized in RIPA buffer (Cell Signaling, #9806), with the addition of PMSF (Thermo Scientific, #36978) and Protease (Sigma-Aldrich, #P8340) inhibitors. Homogenates were incubated in ice for 30 minutes with periodical pipetting and centrifuged at 15.000 x g for 15 minutes. Proteins were quantified by Bradford Assay (Thermo Scientific, #1863028) and equal amounts were loaded for SDS-PAGE and transferred to Immobilon-Fl PVDF membrane (Millipore, IPFL00010). Each membrane was blocked for 1.5 hr at Room Temperature with Odyssey Blocking Buffer (TBS) (Li-Cor, #927-50000). Membranes were incubated with primary antibodies at 4°C overnight, washed, and then blots were incubated for 1.5 hr at Room Temperature with the corresponding secondary antibodies (Li-Cor). After incubation with secondary antibodies, blots were scanned by Odyssey CLx Imaging system (Li-Cor). The band intensity was automatically determined by the accompanying software Image Studio Ver 5.2 (Li-Cor). GAPDH was used as loading controls. Antibodies and dilutions are shown in Table 2.

**Table 2:** Antibodies used for immunofluorescence and western blot.

| **Protein** | **Dilution** | **Manufacturer** | **Use** |
| --- | --- | --- | --- |
| RyR | 1:30 (IHC); 1:40 (WB) | 34C, DSHB | WB, IHC |
| DHPR (CaV1.1) | 1:400 | Invitrogen | WB |
| STIM1 (rabbit) | 1:1000 (WB) 1:100 (IHC) | S6197, Sigma | WB, IHC |
| STIM1 (mouse) | 1:50 | 610954, BD Biosciences | IHC |
| GAPDH | 1:50000 | Thermo Fisher | WB |
| ORAI1 | 1:20 | PA5-26378, Thermo Scientific | IHC |
| SERCA | 1:100 | DSHB | WB |
| Alexa Fluor goat anti-mouse 488 IgG | 1:500 | Invitrogen, A11029 | IHC |
| Alexa Fluor goat-anti rabbit 568 IgG | 1:500 | Invitrogen, A11036 | IHC |

### EDL muscle bundles and immunofluorescence

EDL muscles from 6-month-old, C57BL/6J WT and C3KO male mice were harvested at no treadmill exercise (control), 1-hour post treadmill-exercise, and 6-hour post-treadmill exercise time points and used for immunofluorescent staining, as previously described (20). EDL muscles were excised, pinned at the proximal and distal tendons and fixed in 2% paraformaldehyde at a length slightly stretched past resting length, for two hours at room temperature. After fixation, EDLs were gently separated into small muscle bundles from the distal tendons. Bundles were washed three times in 1x PBS for 5 minutes each wash, permeabilized for 2 hours at room temperature with 2% Triton-X in PBS, then blocked in 10% goat serum with 0.5% Triton-X for 1 hour at room temperature with gentle rocking. Bundles were incubated overnight at 4°C with antibodies recognizing RyR, STIM1 and ORAI1 (Table 2), diluted in blocking solution. After overnight incubation, primary antibodies were removed, bundles were washed three times in 1x PBS for 5 minutes each wash, then incubated with secondary antibodies for two hours at room temperature in the dark, with gentle agitation. Bundles were finally washed in PBS 3 times after secondary antibody incubation, mounted with VECTASHIELD Antifade Mounting Medium (Vector, H-1000) and visualized using a Leica Stellaris 5 Confocal Microscope equipped with a 40x oil-immersion objective (Leica Microsystems Inc, Illinois, USA). Fluorescence intensity profiles across sarcomeres were plotted offline using the FIJI software.

### Electron Microscopy

Electron microscopy was performed in EDL muscles as previously described (20) from WT and C3KO mice at no exercise, 1 and 6 hours post-exercise. Briefly, EDL muscles were dissected from euthanized sedentary and exercised animals, pinned on a Sylgard dish and fixed at room temperature with 3.5% glutaraldehyde in 0.1 M sodium cacodylate (NaCaCo) buffer (pH 7.2). Small bundles of fixed muscle were post-fixed, embedded, stained en-block, and sectioned for EM. For T tubule staining, dissected EDL were fixed in 2% glutaraldehyde in NaCaCo buffer and were post-fixed in a mixture of 2% OsO4 and 0.8% K3Fe(CN)6 for 1–2 h. Ultrathin sections (∼50 nm) were cut using a Leica Ultracut R microtome (Leica Microsystem) with a Diatome diamond knife (Diatome Ltd.) and double-stained with uranyl acetate and lead citrate. Sections were viewed and photographed using a 120 kV JEM-1400 Flash Transmission Electron Microscope (Jeol Ltd., Tokyo, Japan) equipped with CMOS camera (Matataki and TEM Center software Ver. 1.7.22.2684 (Jeol Ltd., Tokyo, Japan)).

### Quantitative Analyses of EM Images

For each different group we analysed 3 mice. All EM quantifications were performed in cross sectioned muscle fibers.

1. The incidence of SR stacks were determined from electron micrographs of nonoverlapping regions randomly collected by counting the number of stacks per area of section (100 μm^2^) as described previously (20, 23). In WT muscles, 4-5 representative fibers were analyzed, while for each of the other groups, 15-20 representative fibers were analyzed, and 5 micrographs at 10,000-12,000× magnification were taken for each fiber.
2. Extensions of the T-tubule network within the I band sarcomere (i.e. total T-tubule) length and T-tubule/SR stack contact length (i.e., length of the association between T-tubule and SR stack membranes) were measured in electron micrographs of nonoverlapping regions randomly collected from muscle fibers after staining of T-tubules with potassium ferrocyanide. For each specimen, 10–15 representative fibers were analyzed, and 5 micrographs at 10,000-12,000× magnification were taken for each fiber.
3. The incidence of T-tubule/SR stack junctions (i.e., CEUs; 100 μm2) were determined from electron micrographs of nonoverlapping regions randomly collected by counting the number of CEUs per area of section (100 μm^2^) as described previously (42). In WT muscles, 4-5 representative fibers were analyzed, while for each of the other samples, 15-20 representative fibers were analyzed, and 5 micrographs at 10,000-12,000× magnification were taken for each fiber.

### Isolation of single FDB muscle fibers

The extraction and analysis of single FDB muscle fibers were conducted to measure electrically-evoked Ca^2+^ transient and SOCE. FDB muscles were extracted from mouse footpads and digested in 1.3 mg/ml collagenase A (Roche Diagnostics, Indianapolis, IN, USA) in Ringers solution containing 146 mM NaCl, 5 mM KCl, 2 mM CaCl_2_, 1 mM MgCl_2_, and 10 mM HEPES (pH 7.4), at 37°C for 75 minutes with gentle rocking. FDB fibers were dissociated through gentle trituration and plated onto glass-bottom dishes until use.

### Ca^2+^ transient measurements

Myoplasmic Ca^2+^ transients were measured as previously described (21, 23). In brief, acutely isolated FDB fibers were treated with 4 µM mag-fluo-4-AM, a moderate-affinity Ca^2+^ indicator, for 20 minutes at room temperature, followed by a washout in a dye-free solution for 20 minutes, supplemented with a skeletal muscle myosin inhibitor, 10 µM N-benzyl-p-toluene sulfonamide (BTS) to minimize cell movement during the stimulation. Loaded FDB fibers were mounted onto the stage of a Nikon Ti2U inverted microscope equipped with a 40x oil immersion objective, and then subjected to a repetitive stimulation protocol with 40 consecutive, 500-ms stimulations at 50-Hz delivered every 2.5 seconds, duty cycle 0.2. The stimulation was performed using an extracellular electrode placed adjacent to the cell of interest. Mag-fluo-4 was excited by 480 ± 15 nm epifluorescent light (Excite epifluorescence illumination system, Nikon Instruments, Melville, NY, USA). The fluorescence emission was detected by a photomultiplier detection system (Photon Technologies Inc, Birmingham, NJ, USA) and collected by a pClamp software. The peak Ca^2+^ transient amplitude was measured at the end of each tetanus and expressed as (Fmax-F_0_)/F_0_ (ΔF/F_0_) using Clampfit software (Molecular Devices, Sunnyvale, CA, USA).

For resting cytosolic Ca^2+^ measurements, FDB fibers were loaded with 5µM Indo-1 AM at room temperature for 30 minutes, followed by 20 minutes washout in dye-free Ringer’s solution. Indo-1 was excited by 365 ± 5nm epifluorescent light and detected at 405nm and 485nm. Ratio of fluorescence emission at 405nm and 485nm (F_405nm_/F_485nm_) was used to determine resting Ca^2+^.

### SOCE measurements

To determine the maximum rate of SOCE in WT and C3KO mice at rest, FDB fibers were treated with a SR store depletion cocktail (30 µM cyclopiazonic acid, and 2 µM thapsigargin in a Ca^2+^-free Ringers solution containing: 146 mM NaCl, 5 mM KCl, 1 mM MgCl_2_, 0.2mM EGTA and 10 mM HEPES, pH 7.4) for 1 hour at 37°C, to facilitate store depletion and SOCE activation. Indo-1 AM was also included in the depletion cocktail for simultaneous dye loading. Following store depletion and dye loading, FDB fibers were washed with Ca^2+^-free Ringer’s solution supplemented with 25 µM BTS for 15 minutes before being imaged. In one subset of experiments, FDB fibers were measured for SOCE activity without depleting the SR Ca^2+^ store, with the fiber treated the same way as above, excluding the SERCA pump inhibitors (cyclopiazonic acid and thapsigargin) in the solution.

Post treatment, store-depleted or non-depleted FDB fibers were immersed in Ca^2+^-free Ringer’s solution with 25 µM BTS. Indo-1 was excited by 365 ± 5nm epifluorescent light and fluorescence intensity detected at 405 nm and 485 nm. Changes in F_405nm_/F_485nm_ ratio were used to monitor Ca^2+^ influx through SOCE, when fibers were first extracellularly perfused for 30 seconds with 30 mM Caffeine to ensure store depletion and then with Ca^2+^ (2 mM) containing Ringer’s solution for 600 seconds. The peak of SOCE (R_SOCE_) was computed as R_SOCE_ = R_max_-R_baseline_ and denoted as Indo-1 Δ Ratio. Analysis was performed using the Clampfit 10.6 software (Molecular Devices, Sunnyvale, CA, USA).

### Statistical Analysis

Statistical analysis and data visualizations were generated in GraphPad Prism 9. SOCE and resting Ca^2+^ data analyzed with unpaired t-test. Fatigue muscle mechanics and Ca^2+^ transient data analyzed with multiple unpaired *Student* t-test. One-way ANOVA with Tukey *post-hoc* test was used for multiple group comparisons. Two-way ANOVA followed by Sidak’s post-hoc testing was used for comparisons of ORAI1 peaks. All graphs are averages with error bars representing SEM. P < 0.05 was considered statistically significant.

### Author Contributions

LWL and ERB conceived and designed the study. KV, RZ, CSHB, and GR performed all experiments. All authors analyzed the data. KV, RZ, ERB, and LWL wrote the manuscript.

## ACKNOWLEDGMENTS

This work was supported by the Coalition to Cure Calpain 3 Foundation (AGR00024098) to ERB and LWL, and by US National Institute of Health (NIH) P50 AR052646 to ERB supporting the Physiological Assessment Core. KV also received support from P50 AR052646 Training Core. We are grateful to Dr. Melissa Spencer for providing the C3KO mice to generate the mice in this study.

